# Biomarker identification for statin sensitivity of cancer cell lines

**DOI:** 10.1101/215756

**Authors:** Vineet K. Raghu, Colin H. Beckwitt, Katsuhiko Warita, Alan Wells, Panayiotis V. Benos, Zoltán N. Oltvai

## Abstract

Statins are potent cholesterol reducing drugs that have been shown to reduce tumor cell proliferation *in vitro* and tumor growth in animal models. Moreover, retrospective human cohort studies demonstrated decreased cancer-specific mortality in patients taking statins. We previously implicated membrane E-cadherin expression as both a marker and mechanism for resistance to atorvastatin-mediated growth suppression of cancer cells; however, a transcriptome-profile-based biomarker signature for statin sensitivity has not yet been reported. Here, we utilized transcriptome data from fourteen NCI-60 cancer cell lines and their statin dose-response data to produce gene expression signatures that identify statin sensitive and resistant cell lines. We experimentally confirmed the validity of the identified biomarker signature in an independent set of cell lines and extended this signature to generate a proposed statin-sensitive subset of tumors listed in the TCGA database. Finally, we predicted drugs that would synergize with statins and found several predicted combination therapies to be experimentally confirmed. The combined bioinformatics-experimental approach described here can be used to generate an initial biomarker sensitivity for statin therapy.

## INTRODUCTION

Despite advances in cancer therapy in past decades, cancer remains the second leading cause of death in the United States [1]. The high cost and length of novel drug development motivates the repurposing of existing drugs, especially since their safety profiles are well-established [2, 3]. This goal, at least in part, can be supported by identifying biomarker sensitivity signatures for existing anticancer therapies, and by predicting drug combinations that would augment the effectiveness of monotherapies.

The HMG-CoA reductase (HMGCR) inhibitors, statins, have been clinically approved for the treatment of hypercholesterolemia for thirty years [4]. Large retrospective studies of statin usage in cancer patients have shown that while statins do not affect cancer incidence [5, 6], their use appears to reduce cancer mortality [7, 8]. These studies have been supported by experimental data that show anti-tumor effects of statins on many cancer cell lines and in some animal models by inducing apoptosis or growth arrest [9–13]. Not all tumor cell lines are sensitive to statins, however, and prospective clinical trials have reported ambiguous outcomes [14]. Thus, a gene expression signature of statin sensitivity would enable researchers and clinicians to focus on predicted sensitive and resistant cell lines, tissues, and patients for further mechanistic and clinical studies. Moreover, those predictions would identify candidate biomarkers and genes that play a role in tumor susceptibility or resistance to statins. Finally, this model may suggest a role for statins in anticancer therapy for patients with predicted statin-sensitive tumors.

Here, we satisfied these aims by utilizing transcriptome data from fourteen NCI-60 cancer cell lines and their sensitivity data to two statin drugs to produce a genetic signature identifying statin sensitive cells. We enriched these data with publicly available gene expression data from the National Cancer Institute (NCI-60) and the Cancer Cell Line Encyclopedia (CCLE), and with biomarker discovery algorithms that can distinguish direct from indirect interactions in large datasets. We experimentally confirmed the validity of the identified biomarker signature in an independent set of cell lines, showed that a subset of TCGA tumors are predicted to display statin sensitivity, and demonstrated the biological viability of the predicted signature. The combined bioinformatics-experimental approach described here can be used to generate biomarker sensitivity signatures for anticancer therapies and generate hypotheses of mechanism of action of drug sensitivity and resistance.

## MATERIAL AND METHODS

### Cell culture, statin treatment and cell proliferation assay

We selected seven pairs of cell lines from the NCI-60 cancer cell panel. These cell lines represent seven different major solid tumor types. For each site, we selected one cell line with low and one with high protein synthesis rate, as previously reported [15]. The selected cell lines - colon cancer (HCT-116, KM-12), ovarian cancer (IGROV1, OVCAR3), breast cancer (HS-578T, T47D), lung cancer (HOP-92, NCI-H322M), prostate cancer (PC-3, DU-145), melanoma (SK-MEL-5, MDA-MB-435), and brain cancer (SF-295, SF-539) - were cultured in RPMI 1640 medium (Life Technologies, Grand Island, NY), supplemented with 10% heat-inactivated fetal bovine serum (HI-FBS, Life Technologies) and 0.5% penicillin/streptomycin (Life Technologies) at 37°C with 5% CO_2_.

Atorvastatin (Sigma-Aldrich, St. Louis, MO) and Rosuvastatin (Santa Cruz Biotechnology, Santa Cruz, CA) were dissolved in dimethyl sulfoxide (DMSO, Sigma-Aldrich; final concentration of 0.1% in RPMI 1640 medium) at a final concentration of 10 µM. The cells were seeded in 6-well plates at a density of 1×10^5^ cells/ml and incubated overnight prior to treatment with 10 µM Atorvastatin, 10 µM Rosuvastatin, or 0.1% DMSO (DMSO served as solvent control). Three independent experiments were performed. Cell proliferation was quantified at 2, 4, and 6 days by direct cell counting with Scepter^TM^ Handheld Automated Cell Counter (EMD Millipore, Billerica, MA) using Scepter^TM^ Tips‒60 µm sensor (EMD Millipore).

Description of the IC_50_ determination, Immunofluorescence microscopy and Computational Models can be found in the Supplementary Materials.

## RESULTS

### Biomarker identification for statin sensitivity of cancer cell lines using temporal statin growth inhibition data in fourteen cancer cell lines

Previous experiments have demonstrated that statins, including atorvastatin (Lipitor), inhibit the growth of a subset of NCI-60 cancer cell lines, and that a subset of statins show similar half-maximal inhibitory concentration (IC_50_) values [16]. To test if the chemical properties of statins affect their inhibition of tumor cells, we cultured fourteen cancer cell lines from the NCI-60 collection, derived from seven organ types, in standard growth medium in the presence of a lipophilic (10 µM atorvastatin) or hydrophilic (10 µM rosuvastatin) statin. Importantly, in enzyme assays these statins have similar affinities for their enzyme target, HMG-CoA reductase (HMGCR), with IC_50_ values of 8.2nM and 5.4nM for atorvastatin and rosuvastatin, respectively [17]. We previously demonstrated that atorvastatin affected the growth of these cancer cell lines differentially; some cell lines displayed full or partial growth inhibition while others were completely insensitive to atorvastatin treatment [15]. In the current study, we have found that all the atorvastatin sensitive cell lines except HOP-92 are partially or fully resistant to rosuvastatin at the tested drug concentrations (Fig. 1).

**Figure 1.**
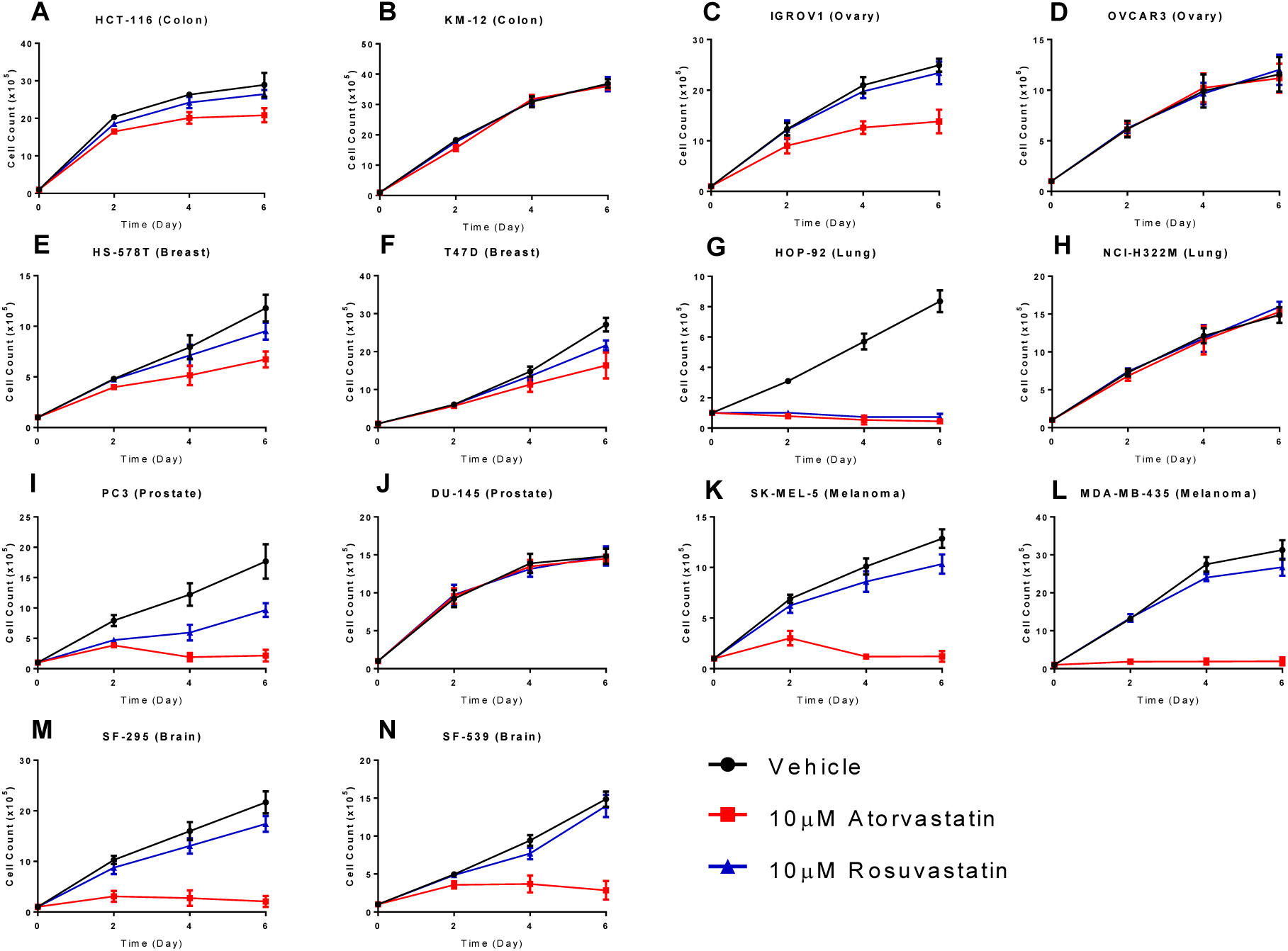
Growth rate of atorvastatin and rosuvastatin treated NCI-60 cancer cell lines. Colon cancer- (**A**. HCT-116 and **B**. KM-12), ovarian cancer- (**C**. IGROV1 and **D**. OVCAR3), breast cancer- (**E**. HS578T and **F.** T47D), lung cancer- (**G.** HOP-92 and **H.** NCI-H322M), prostate cancer- (**I.** PC-3 and **J**. DU-145), melanoma- (**K**. SK-MEL-5 and **L**. MDA-MB-435), and brain cancer- (**M**. SF-295 and **N**. SF-539) cell lines from the NCI-60 cancer cell line collection were treated with 10 µM atorvastatin (pink line), 10 µM rosuvastatin (blue line), or DMSO vehicle control (black line) and cell growth was quantified at 2, 4, and 6 days through direct cell counting. The results are representative of three independent experiments. Error bars represent the standard deviation (n=3).

We utilized gene expression data from these fourteen NCI-60 cancer cell lines [16, 18] and statin dose-response data (see Fig. 1) to produce a genetic signature distinguishing between atorvastatin and rosuvastatin sensitive and resistant cell lines. Specifically, our training set consisted of baseline gene expression data for the fourteen (non-treated) NCI-60 cell lines, and a measure of statin sensitivity as our response variable. We computed this response variable over a growth period of six days utilizing atorvastatin (ato), rosuvastatin (rosu), ato + rosu, or vehicle treatment on each cell line. From these data, dose-response curves were formed, depicting the number of (proliferating) cells at each time point. Using these curves, we found the ratio of the area under the dose response curve for the control and the area under the dose response curve for the treatment group, and used this number as a final response variable for each cell line.

To determine a genetic signature of statin sensitivity, the variables that were found to be associated to the sensitivity score measure from our graphical modeling methods were selected as important features. Specifically, we used three different methods for feature selection in this analysis. First, we used the “Atorva MGM” approach, which simply used the MGM model to produce an undirected graphical network from the NCI-60 data, and used all the genes adjacent to the statin sensitivity variable in the network as features. Nineteen genes were selected by this approach (Table 1) (Fig. S1a). To determine the reliability of the selected genes, we employed a leave-one-out cross validation method using an SVM regression model, and we obtained a cross validation error rate of 0.0082 for this set of features.

**Table 1.** Selected biomarker for each identification method. Overlapping genes from all three methods are shown in red, and overlapping genes from two methods are shown in bold.

| Atorva-MGM | Combined-MGM | Atorva-TReCS |
| --- | --- | --- |
| **GABRA3** | **GABRA3** | **GABRA3** |
| **TM4SF18** | **TM4SF18** | **TM4SF18** |
| **ATP8B1** |  | **ATP8B1** |
| **CKAP4** |  | **CKAP4** |
| **IL13RA2** | **IL13RA2** |  |
| **KRT7** |  | **KRT7** |
| **LAMA4** |  | **LAMA4** |
| **MACC1** |  | **MACC1** |
| **MISP** |  | **MISP** |
| **MMP14** |  | **MMP14** |
| **NFKBIZ** |  | **NFKBIZ** |
| **SCNN1A** |  | **SCNN1A** |
| **SLC16A4** |  | **SLC16A4** |
| **SLC27A2** |  | **SLC27A2** |
| **SLC6A8** | **SLC6A8** |  |
| **SPINT2** |  | **SPINT2** |
| **TNFRSF10D** | **TNFRSF10D** |  |
| **HIST1H1A** | **ANPEP** | **CD70** |
| **SRPX** | **GFPT2** | **CTSK** |
|  | **PTGFRN** | **IFITM2** |
|  | **RGS4** |  |

Our second approach was the “Combined” approach. Here, we used the MGM selected features from rosuvastatin on the NCI-60 dataset with the sensitivity scores from the rosuvastatin experiments. We combined them with the MGM selected features from the atorvastatin cell lines, and applied T-ReCS on the NCI-60 dataset to produce clusters of relatively overexpressed or suppressed genes predictive of statin sensitivity. Following this step, we searched for those genes from the rosuvastatin MGM selected features that were in any cluster from the atorvastatin features selected. These were then utilized as the genetic signature. In total, nine genes were selected using this approach (Table 1)(Fig. S1b), but the cross validation error rate was higher (0.1967).

Our final approach was the Atorva_TReCS approach. Here, we used the MGM features for the atorvastatin data as the original features for T-ReCS. Then, we ran our T-ReCS algorithm to produce clusters on these features. We then choose the final selected features for prediction, by simply taking a single gene from each cluster that had the highest correlation to the statin sensitivity variable. In total, seventeen genes were selected using this approach (Table 1) (Fig. S1c) (as some genes from the original MGM selection appeared in the same cluster), with a cross validation error rate of 0.0285.

For each of these approaches, we used the corresponding gene expression signature to train a predictive linear regression model (an SVM regression model trained via MATLAB’s “fitrsvm” library) on the NCI-60 gene expression data. To validate the predictive power of this model, we obtained gene expression data from the Broad Institute’s Cancer Cell Line Encyclopedia (CCLE) [19], and this data was used to predict an Atorvastatin sensitivity score for each of the cell lines in the CCLE database. We utilized the predictions from each of the three genetic signature selection approaches in the following manner. First, using the genes selected from each of the methods the top ten statin resistant cell lines were predicted using the model trained on the NCI-60 database. Then, each of these cell line sets were combined and duplicates were removed resulting in 26 resistant cell lines. The same approach was employed on the set of 23 predicted statin sensitive cell lines. It is evident that the predictions by the three methods yielded a partial overlap for both the most statin sensitive and statin resistant cell lines (see Supplementary Table 5 [excel sheet] in the Supplement).

### Experimental validation of biomarker predictions

Next, we assessed the fidelity of the predicted biomarkers by experimentally testing the statin sensitivity of yet untested cell lines. To reduce the number of cell lines to be experimentally tested, we clustered the predicted 23 statin sensitive and 26 statin resistant cell lines based on their whole transcriptome profiles, since cells displaying highly similar transcriptome profiles are likely to have similar biologic behavior. We performed principal component analysis on the full gene expression data for the predicted 26 statin resistant cell lines, and the cell lines were clustered according to their first two principal components using a *k*-means clustering with *k* = 10 (Fig. S2a). The same approach was employed on the set of 23 predicted statin sensitive cell lines (Fig. S2b).

To test the atorvastatin sensitivity of representative cell lines from the predicted sensitive and resistant groups, we selected the NCI-H2170 (lung cancer-derived) and BT-474 (breast cancer-derived) predicted statin resistant; and the SK-MES-1 (lung cancer-derived) and SK-MEL-24 (melanoma-derived) predicted statin sensitive cell lines. To determine the IC_50_ of atorvastatin, we treated the cell lines with half log doses of atorvastatin, between 100nM and 100uM. As shown in Figure 2, cell lines predicted to be sensitive (SK-MES-1 and SK-MEL-24) were more sensitive to atorvastatin than the cell lines predicted to be resistant (NCI-H2170 and BT-474) to it (Fig. 2a, b), confirming the fidelity of the predicted statin-sensitivity gene expression signature.

**Figure 2.**
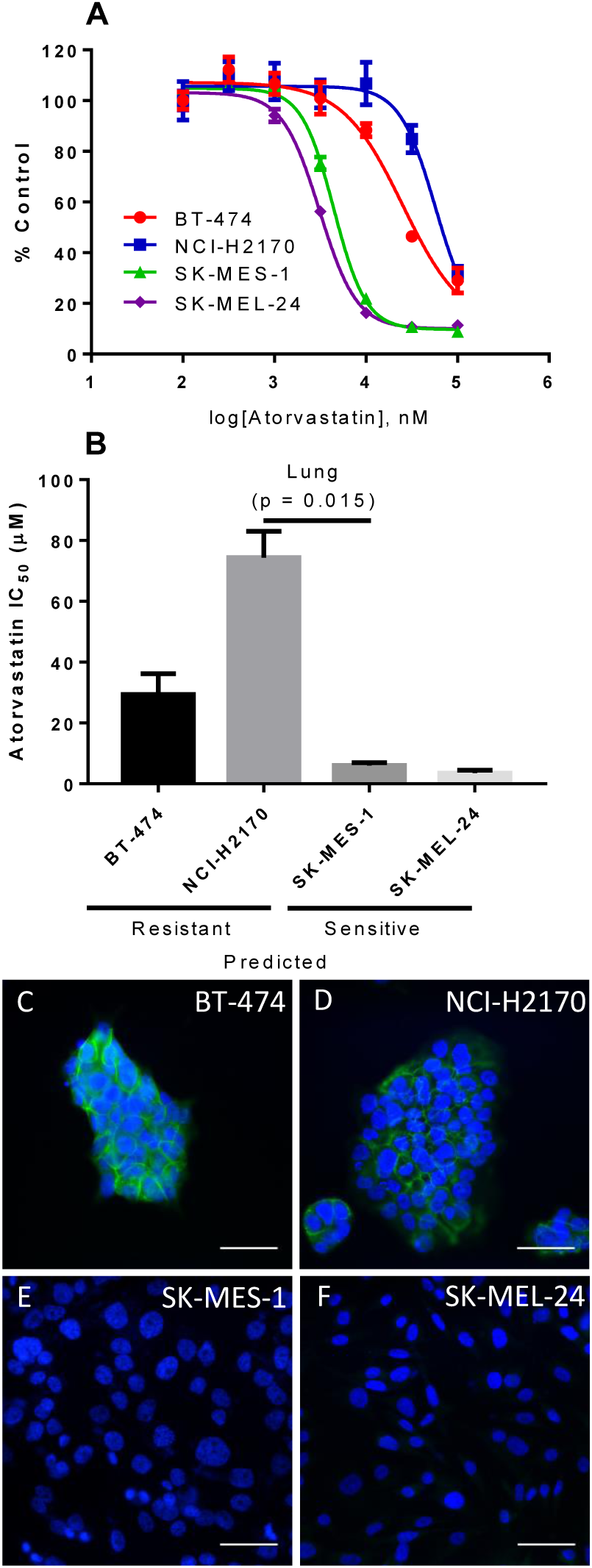
Empirical statin sensitivity data correlates with predictions and membrane E-cadherin expression. (A.) Dose response curves for Atorvastatin in predicted resistant (BT-474 and NCI-H2170) and predicted sensitive (SK-MES-1 and SK-MEL-24) cell lines. (B) Predicted resistant cell lines exhibited higher IC_50_ values to atorvastatin than predicted sensitive cell lines. Data are representative of three independent experiments. All experiments were carried out in triplicate (N=3). Error bars represent the standard error of the mean. [19] Atorvastatin resistant cell lines BT-474 (C) and NCI-H2170 (D) display strong membrane E-cadherin (green) staining. In contrast, atorvastatin sensitive cell lines SK-MES-1 (E) and SK-MEL-24 (F) exhibited no E-cadherin staining. DAPI (blue) was used to counterstain the cell nuclei. Scale bar = 50µm.

Statin resistance correlates with cell membrane E-cadherin expression in cancer cell lines [15]. To determine the expression of E-cadherin in BT-474, NCI-H2170, SK-MES-1, and SK-MEL-24 cell lines we immunostained these cell lines for E-cadherin, as previously described [15]. We find that statin resistant cells with high mRNA for E-cad (BT-474 and NCI-H2170) have high membrane E-cadherin expression. In contrast, statin-sensitive cells with low mRNA for E-cad (SK-MES-1 and SK-MEL-24) have no detectable E-cadherin staining (Fig. 2c-f). The uncovered E-cadherin expression pattern agrees with E-cadherin mRNA expression data deposited for these cell lines in Oncomine [20] and demonstrates that membrane E-cadherin is a marker of statin-resistant cells.

### Determination of potential combination statin therapies

To determine if there was potential for combination therapies with statins across cell lines, we analyzed publicly available pharmacological profiles from the Cancer Cell Line Encyclopedia (CCLE) and the Sanger Center Genomics of Drug Sensitivity project. The CCLE data profiled 24 molecules across 504 cell lines [19], and the Sanger Center profiled 250 molecules across 549 cell lines [21]. Cell line sensitivity measurements for each molecule on each profiled cell line was computed using the IC_50_ value and the area under the dose response curve (AUC) for the CCLE data and the Sanger center data, respectively. These sensitivity measurements were compared to predicted statin sensitivity on only those cell lines appearing in both the CCLE baseline gene expression dataset as well as the pharmacological profiling datasets, and correlation scores were produced (<u>Table 2</u>). Further discussion on these results can be found in the Supplement.

**Table 2.** Molecules with similar pharmacological profiles to predicted statin sensitivity across cell lines. Pharmacological profiles from CCLE and Sanger Center drug sensitivity data were used to predict combination therapy with statins. Molecules with positive correlation and FDR Q-Value < .05 were included. For CCLE profiles, IC_50_ was used as a proxy for sensitivity, and for Sanger center profiles, the area under the dose-response curve (AUC) was used.

| Drug Name | Target | Correlation | P Value |
| --- | --- | --- | --- |
| **Dabrafenib** | BRAF | 0.456976009 | 2.38E-18 |
| **PLX-4720** | BRAF | 0.376159659 | 4.97E-13 |
| **(5Z)-7-Oxozeaenol** | TAK1 | 0.376014976 | 8.61E-13 |
| **SB590885** | BRAF | 0.340372121 | 4.32E-09 |
| **TW 37** | BCL2, BCL-XL, MCL1 | 0.27305505 | 1.89E-06 |
| **Piperlongumine** | NF-κB | 0.226445462 | 0.000199099 |
| **Cisplatin** | DNA crosslinker | 0.233847531 | 0.000250235 |
| **Lestauritinib** | FLT3, JAK2, NTRK1, NTRK2, NTRK3 | 0.218300548 | 0.000905096 |
| **Elesclomol** | HSP90 | 0.206861021 | 0.002058482 |
| **Refametinib** | MEK1, MEK2 | 0.183970201 | 0.004239963 |
| **Bleomycin** | dsDNA break induction | 0.183339449 | 0.004370418 |
| **Temsirolimus** | MTOR | 0.182232337 | 0.01021699 |
| **Nutlin-3a (-)** | MDM2 | 0.176424024 | 0.010924768 |
| **Selumetinib** | MEK1, MEK2 | 0.165658468 | 0.010924768 |
| **AZ628** | BRAF | 0.296949736 | 0.011550955 |
| **PD0325901** | MEK1, MEK2 | 0.173101761 | 0.011550955 |
| **Serdemetan** | MDM2 | 0.164726002 | 0.011550955 |
| **Trametinib** | MEK1, MEK2 | 0.166312205 | 0.011550955 |
| **Docetaxel** | Microtubule stabilizer | 0.169092858 | 0.013420485 |
| **BX796** | TBK1, PDK1 (PDPK1), IKK, AURKB, AURKC | 0.167710541 | 0.013718152 |
| **Embelin** | XIAP | 0.157305764 | 0.016974867 |
| **FH535** | PPARγ, PPARΔ | 0.154672573 | 0.02034349 |
| **YK-4-279** | RNA helicase A | 0.158950401 | 0.026108609 |
| **Rucaparib** | PARP1, PARP2 | 0.147699505 | 0.027477555 |
| **CI-1040** | MEK1, MEK2 | 0.152967932 | 0.032836113 |
| **SN-38** | TOP1 | 0.140036378 | 0.040080871 |
| **CHIR-99021** | GSK3A, GSK3B | 0.135343422 | 0.041404418 |

## DISCUSSION

Despite many advances in the treatment of local and advanced cancers, metastatic disease still exhibits high mortality and its management remains complex. The genomic instability of cancers that have undergone or are prone to metastasis promotes the development of drug resistance, hindering mono-therapy for these advanced cancers. As such, combination therapies are needed to combat tumor progression by preventing cellular compensation to a single drug treatment. In order to thoughtfully design new drug combinations, models that incorporate genomic information and provide drug sensitivity predictions are invaluable.

In this work, we have utilized several feature selection approaches to identify a predictive genetic signature of statin sensitivity from RNA-Seq data from TCGA. We applied this signature to several cancer cell lines from the NCI-60 database and externally validated the sensitivity predictions, confirming all four of our experimental predictions. We then applied our predictive signature to the TCGA-Breast Invasive Carcinoma project and found a subset of tumor samples predicted to be sensitive to statins (Supplementary Figures 3A-3I, and we further validated our predictive signature through an ontological analysis for relevance of the selected genes (Supplementary Tables 1-3).

Since the advent of high-throughput gene sequencing technology, genetic biomarker discovery on large datasets has been extensively studied [22, 23]. Several methods have been developed for biomarker identification to predict sensitivity to anticancer drugs. Most of these methods use penalized regression techniques (Lasso, Elastic Net, Ridge, etc.) along with unsupervised approaches (clustering) on gene expression measurements to directly predict anticancer drug sensitivity across cell lines [24–27]. Though these penalized regression techniques identify relevant genes and are suitable for many prediction tasks, they can be insufficient to identify a mechanistic biomarker signature. Signature identification requires separating indirect associations from direct associations, which is difficult through a single penalized regression alone, and high degrees of multi-colinearity in gene expression data further complicate these searches. However, real biological signature identification has much greater impact than a predictive model alone, as a signature has the potential to provide mechanistic insight alongside accurate predictions. Here we have utilized a signature identification methodology that is able to avoid these issues through the framework of graphical modeling and clustering. Further, we have increased the stability of our results to small changes in the data by incorporating the results from multiple analysis methods. We have demonstrated that this paradigm for signature identification produces viable biological results along with accurate predictions.

However, our approach is limited by the data upon which it was built. First, the vast majority of available large datasets (e.g., TCGA) are derived from primary tumors; this skews the analysis towards relatively well differentiated (epithelial, E-cadherin-positive) cells that are not epidemiologically linked to reduction in mortality from cancer, which is more related to disseminated/ metastatic disease (usually mesenchymal, E-cadherin low/absent). Thus, the tumor selection is not enriched in the tumor cells that are postulated as the direct targets for statins. Second, since the gene expression data are derived from whole tumors, tumor heterogeneity is expected to further obscure the statin sensitivity/resistance signal. Third, gene expression data is inherently limited by high dimensionality (i.e., few cell lines per tumor type compared to the large number of genes/variables), which reduces statistical power. Moreover, predicted sensitivity genes were not examined at the genetic level to determine whether altered expression influences statin efficacy. Finally, predicted synergistic drug combinations with statins were not tested experimentally in these cell lines, although some have been already confirmed in the literature. These limitations are outside of the scope of the current work and will be addressed in subsequent studies.

In summary, we have developed a combined algorithm-experiment blueprint that is able to generate biomarker sensitivity signatures for anticancer therapies, as exemplified here by identifying an initial biomarker signature for statin sensitivity in human cancer cell lines and tumors. This signature was validated both in terms of biological sensibility and by experimental validation, and it confirms many recent physiological effects of statins as shown through our ontological analyses. Future studies will be aimed at experimentally testing many more cancer cell lines for their statin sensitivity and better characterizing the genetic factors that impact cancer cell response to statin mono- and combination therapy and experimental verification of these findings in sophisticated *ex vivo* and *in vivo* models of metastatic cancer.

## Data Availability

All data generated or analyzed during this study are included in this published article (and its Supplementary Information files).

## ACKNOWLEDGEMENT

We thank J.R. Chaillet for comment on the manuscript. This work was supported by the National Institutes of Health [TR000496 to A.W., T32EB001026 to C.B., T32CA082084 to V.R., R01LM012087 and U01HL137159 to P.V.B.], a Veterans Administration Merit grant to AW, and by the Japan Society for the Promotion of Science KAKENHI grants [JP26890019 and JP16K18439 to KW].

## SUPPLEMENTARY DATA

Supplementary Data are available online.

## References

[1] R.L. Siegel, K.D. Miller, A. Jemal, Cancer statistics, 2016, CA: a cancer journal for clinicians, 66 (2016) 7–30.

[2] M. Allison, NCATS launches drug repurposing program, Nature biotechnology, 30 (2012) 571–572.

[3] S.N. Deftereos, C. Andronis, E.J. Friedla, A. Persidis, A. Persidis, Drug repurposing and adverse event prediction using high-throughput literature analysis, Wiley interdisciplinary reviews. Systems biology and medicine, 3 (2011) 323–334.

[4] A. Endo, A historical perspective on the discovery of statins, Proceedings of the Japan Academy. Series B, Physical and biological sciences, 86 (2010) 484–493.

[5] J. Haukka, R. Sankila, T. Klaukka, J. Lonnqvist, L. Niskanen, A. Tanskanen, K. Wahlbeck, J. Tiihonen, Incidence of cancer and statin usage--record linkage study, International journal of cancer, 126 (2010) 279–284.

[6] J. Shepherd, G.J. Blauw, M.B. Murphy, E.L. Bollen, B.M. Buckley, S.M. Cobbe, I. Ford, A. Gaw, M. Hyland, J.W. Jukema, A.M. Kamper, P.W. Macfarlane, A.E. Meinders, J. Norrie, C.J. Packard, I.J. Perry, D.J. Stott, B.J. Sweeney, C. Twomey, R.G. Westendorp, Pravastatin in elderly individuals at risk of vascular disease (PROSPER): a randomised controlled trial, Lancet (London, England), 360 (2002) 1623–1630.

[7] S.F. Nielsen, B.G. Nordestgaard, S.E. Bojesen, Statin use and reduced cancer-related mortality, The New England journal of medicine, 367 (2012) 1792–1802.

[8] A. Wang, A.K. Aragaki, J.Y. Tang, A.W. Kurian, J.E. Manson, R.T. Chlebowski, M. Simon, P. Desai, S. Wassertheil-Smoller, S. Liu, S. Kritchevsky, H.A. Wakelee, M.L. Stefanick, Statin use and all-cancer survival: prospective results from the Women’s Health Initiative, British journal of cancer, 115 (2016) 129–135.

[9] M.J. Campbell, L.J. Esserman, Y. Zhou, M. Shoemaker, M. Lobo, E. Borman, F. Baehner, A.S. Kumar, K. Adduci, C. Marx, E.F. Petricoin, L.A. Liotta, M. Winters, S. Benz, C.C. Benz, Breast cancer growth prevention by statins, Cancer research, 66 (2006) 8707–8714.

[10] A. Hoque, H. Chen, X.C. Xu, Statin induces apoptosis and cell growth arrest in prostate cancer cells, Cancer epidemiology, biomarkers & prevention: a publication of the American Association for Cancer Research, cosponsored by the American Society of Preventive Oncology, 17 (2008) 88–94.

[11] C. Denoyelle, M. Vasse, M. Korner, Z. Mishal, F. Ganne, J.P. Vannier, J. Soria, C. Soria, Cerivastatin, an inhibitor of HMG-CoA reductase, inhibits the signaling pathways involved in the invasiveness and metastatic properties of highly invasive breast cancer cell lines: an in vitro study, Carcinogenesis, 22 (2001) 1139–1148.

[12] H. Gbelcova, M. Lenicek, J. Zelenka, Z. Knejzlik, G. Dvorakova, M. Zadinova, P. Pouckova, M. Kudla, P. Balaz, T. Ruml, L. Vitek, Differences in antitumor effects of various statins on human pancreatic cancer, International journal of cancer, 122 (2008) 1214–1221.

[13] D.G. Menter, V.P. Ramsauer, S. Harirforoosh, K. Chakraborty, P. Yang, L. Hsi, R.A. Newman, K. Krishnan, Differential effects of pravastatin and simvastatin on the growth of tumor cells from different organ sites, PloS one, 6 (2011) e28813.

[14] P. Gazzerro, M.C. Proto, G. Gangemi, A.M. Malfitano, E. Ciaglia, S. Pisanti, A. Santoro, C. Laezza, M. Bifulco, Pharmacological actions of statins: a critical appraisal in the management of cancer, Pharmacological reviews, 64 (2012) 102–146.

[15] K. Warita, T. Warita, C.H. Beckwitt, M.E. Schurdak, A. Vazquez, A. Wells, Z.N. Oltvai, Statin-induced mevalonate pathway inhibition attenuates the growth of mesenchymal-like cancer cells that lack functional E-cadherin mediated cell cohesion, Sci Rep, 4 (2014) 7593.

[16] R.H. Shoemaker, The NCI60 human tumour cell line anticancer drug screen, Nature reviews. Cancer, 6 (2006) 813–823.

[17] M.K. Wendt, M.A. Taylor, B.J. Schiemann, W.P. Schiemann, Down-regulation of epithelial cadherin is required to initiate metastatic outgrowth of breast cancer, Molecular biology of the cell, 22 (2011) 2423–2435.

[18] U.T. Shankavaram, S. Varma, D. Kane, M. Sunshine, K.K. Chary, W.C. Reinhold, Y. Pommier, J.N. Weinstein, CellMiner: a relational database and query tool for the NCI-60 cancer cell lines, BMC genomics, 10 (2009) 277.

[19] J. Barretina, G. Caponigro, N. Stransky, K. Venkatesan, A.A. Margolin, S. Kim, C.J. Wilson, J. Lehar, G.V. Kryukov, D. Sonkin, A. Reddy, M. Liu, L. Murray, M.F. Berger, J.E. Monahan, P. Morais, J. Meltzer, A. Korejwa, J. Jane-Valbuena, F.A. Mapa, J. Thibault, E. Bric-Furlong, P. Raman, A. Shipway, I.H. Engels, J. Cheng, G.K. Yu, J. Yu, P. Aspesi, Jr., M. de Silva, K. Jagtap, M.D. Jones, L. Wang, C. Hatton, E. Palescandolo, S. Gupta, S. Mahan, C. Sougnez, R.C. Onofrio, T. Liefeld, L. MacConaill, W. Winckler, M. Reich, N. Li, J.P. Mesirov, S.B. Gabriel, G. Getz, K. Ardlie, V. Chan, V.E. Myer, B.L. Weber, J. Porter, M. Warmuth, P. Finan, J.L. Harris, M. Meyerson, T.R. Golub, M.P. Morrissey, W.R. Sellers, R. Schlegel, L.A. Garraway, The Cancer Cell Line Encyclopedia enables predictive modelling of anticancer drug sensitivity, Nature, 483 (2012) 603–607.

[20] D.R. Rhodes, S. Kalyana-Sundaram, V. Mahavisno, R. Varambally, J. Yu, B.B. Briggs, T.R. Barrette, M.J. Anstet, C. Kincead-Beal, P. Kulkarni, S. Varambally, D. Ghosh, A.M. Chinnaiyan, Oncomine 3.0: genes, pathways, and networks in a collection of 18,000 cancer gene expression profiles, Neoplasia (New York, N.Y.), 9 (2007) 166–180.

[21] M.J. Garnett, E.J. Edelman, S.J. Heidorn, C.D. Greenman, A. Dastur, K.W. Lau, P. Greninger, I.R. Thompson, X. Luo, J. Soares, Q. Liu, F. Iorio, D. Surdez, L. Chen, R.J. Milano, G.R. Bignell, A.T. Tam, H. Davies, J.A. Stevenson, S. Barthorpe, S.R. Lutz, F. Kogera, K. Lawrence, A. McLaren-Douglas, X. Mitropoulos, T. Mironenko, H. Thi, L. Richardson, W. Zhou, F. Jewitt, T. Zhang, P. O’Brien, J.L. Boisvert, S. Price, W. Hur, W. Yang, X. Deng, A. Butler, H.G. Choi, J.W. Chang, J. Baselga, I. Stamenkovic, J.A. Engelman, S.V. Sharma, O. Delattre, J. Saez-Rodriguez, N.S. Gray, J. Settleman, P.A. Futreal, D.A. Haber, M.R. Stratton, S. Ramaswamy, U. McDermott, C.H. Benes, Systematic identification of genomic markers of drug sensitivity in cancer cells, Nature, 483 (2012) 570–575.

[22] M.J. Garnett, & McDermott, U., The evolving role of cancer cell line-based screens to define the impact of cancer genomes on drug response, Current opinion in genetics & development, 24 (2014) 114–119.

[23] F. Chibon, Cancer gene expression signatures-the rise and fall?, European journal of cancer, (2013) 2000–2009.

[24] P. Geeleher, Cox, N.J., Huang, R.S., Clinical drug response can be predicted using baseline gene expression levels and in vitro drug sensitivity in cell lines, Genome biology, 15 (2014).

[25] G. Wildey, Chen, Y., Lent, I., Stetson, L., Pink, J., Barnholtz-Sloan, J.S., Dowlati, A., Pharmacogenomic approach to identify drug sensitivity in small-cell lung cancer, PloS one, 9 (2014).

[26] Y. Sun, Zhang, W., Chen, Y., Ma, Q., Wei, J., Liu, Q., Identifying anti-cancer drug response related genes using an integrative analysis of transcriptomic and genomic variations with cell line-based drug perturbations, Oncotarget, 7 (2016).

[27] M.J. Garnett, Edelman, E.J., Heidorn, S.J., Greenman, C.D., Dastur, A., Lau, K.W., Liu, Q., et al., Systematic identification of genomic markers of drug sensitivity in cancer cells, Nature, 483 (2012) 570–575.

